# Selective volume illumination microscopy offers synchronous volumetric imaging with high contrast

**DOI:** 10.1101/403303

**Authors:** Thai V. Truong, Daniel B. Holland, Sara Madaan, Andrey Andreev, Josh V. Troll, Daniel E. S. Koo, Kevin Keomanee-Dizon, Margaret J. McFall-Ngai, Scott E. Fraser

## Abstract

Light field microscopy provides an efficient means to collect 3D images in a single acquisition, as its plenoptic detection captures an extended image volume in one snapshot. The ability of light field microscopy to simultaneously capture image data from a volume of interest, such as a functioning brain or a beating heart, is compromised by inadequate contrast and effective resolution, due, in large part, to light scattering by the tissue. Surprisingly, a major contribution to the image degradation is the signal scattered into the volume of interest by the typical wide-field illumination that excites the sample region outside the volume of interest. Here, we minimize this degradation by employing selective volume illumination, using a modified light sheet approach to illuminate preferentially the volume of interest. This minimizes the unavoidable background generated when extraneous regions of the sample are illuminated, dramatically enhancing the contrast and effective resolution of the captured and reconstructed images. Light Field Selective Volume Illumination Microscopy (LF-SVIM, SVIM for short) dramatically improves the performance of light field microscopy, and offers an unprecedented combination of synchronous z-depth coverage, lateral and axial resolution, and imaging speed.

---

Many biological processes take place over 3-dimensional (3D) spaces extending over hundreds of microns. Studying such dynamic processes across these mesoscopic-scales requires volumetric imaging tools that offer cellular resolution and 3D sectioning, combined with sufficient speed to faithfully capture the cellular dynamics. Typically, such volumetric imaging is accomplished using optical sectioning approaches, in which optical signal is collected one point (confocal), one line (line-confocal), or one plane (light sheet microscopy) at a time; these asynchronously acquired 2D-image planes are then assembled to create a reconstruction of the 3D region of interest ^1,2^. Live specimens can change in shape or physiology on time scales as fast as milliseconds; thus, to avoid distortions, the complete set of 2D images must be collected over similar times, so that they can be assembled into 4-dimensional (4D; x, y, z, time) renderings. However, for any approach in which optical sections are collected in a sequential manner, it is inescapable that different parts of the sample are sampled at different times. Because dynamic information can be lost or distorted by conventional optical sectioning methods, there is an unmet need for a synchronous volumetric live imaging approach.

Light field microscopy (LFM; Fig. 1a) offers an imaging modality with the ability to capture all points in a volumetric image simultaneously. The 3D light field coming from the sample space is recorded on a single 2D camera by positioning a lenslet array in place of the camera at the image plane, and moving the camera one focal length behind the lenslets; this plenoptic detection permits a single 2D camera image to capture information from the 3D volume ^3–5^. Computational reconstruction is used to solve the inverse problem, reconstructing from the recorded 2D image a 3D volumetric image “stack” over an extended z-depth of hundreds of microns, with a trade-off in reduced resolution. When performing LFM in biological tissues, the scattering of light by the tissue cannot be rejected by a confocal aperture as is done in confocal microscopy; this produces a significant signal background which makes solving the inverse problem more challenging and reduces the quality of the reconstructed 3D image. This loss of performance becomes critical when wide-field illumination of the specimen is used, as is done in conventional LFM, since the camera captures not only light coming from the volume of interest, but also a large amount of light scattered from regions far out of the volume of interest (Fig. 1b). This extraneous background captured by the camera reduces the contrast and effective resolution of the final reconstructed images, dramatically limiting the capability to generate useful 4D images in biological tissues.

**Figure 1:**
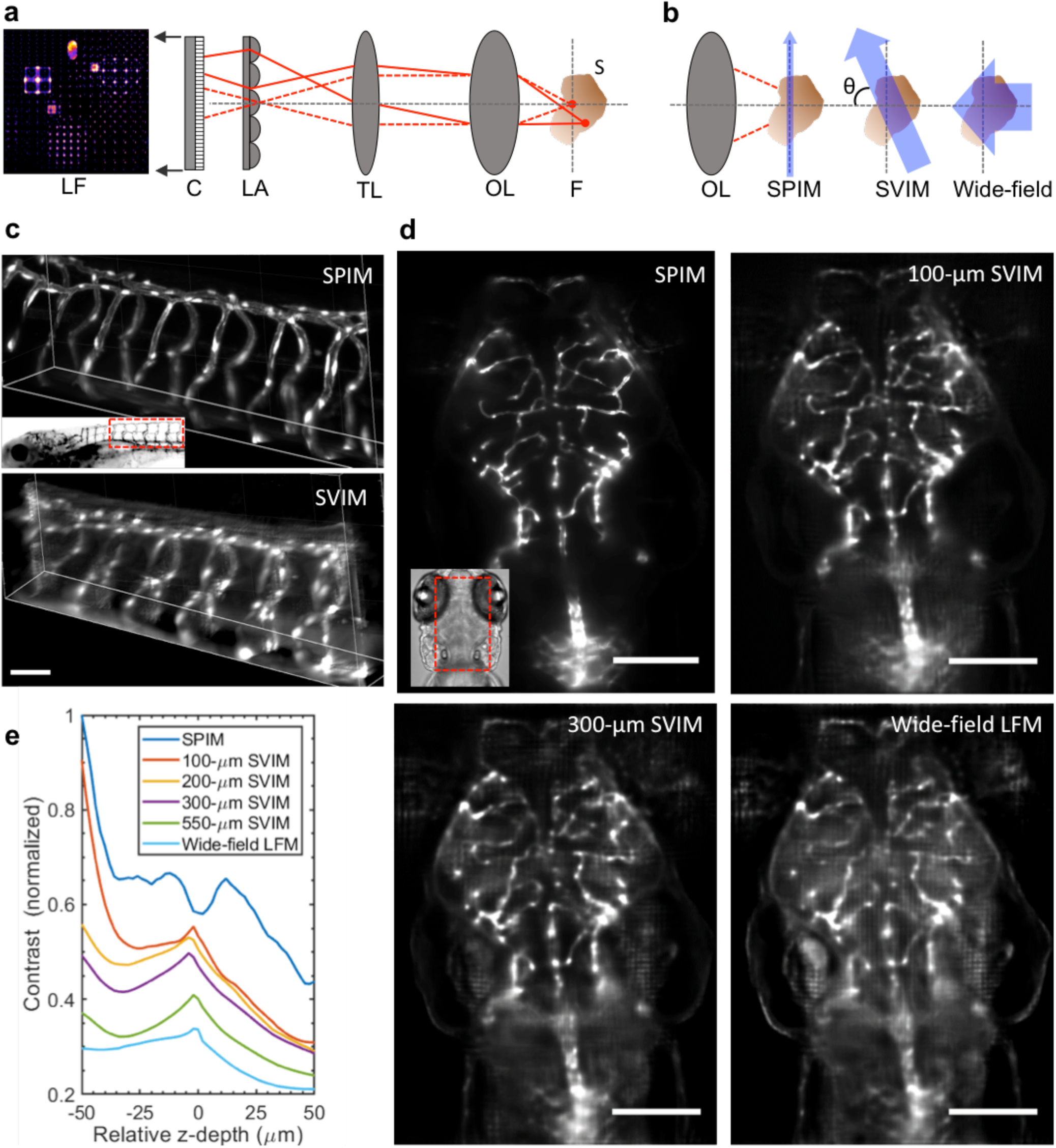
Selective Volume Illumination Microscopy (SVIM) offers improved Light Field Microscopy (LFM) performance for the synchronous imaging of a 3D sample. **(a)** Conventional light field microscopy extends a conventional microscope, which produces a magnified image of the sample (S) from the native focal plane (F) to the image plane (IP) using an objective lens (OL) and tube lens (TL). To synchronously capture information at z-positions above and below F, a micro-lens array (LA) is placed at the IP, encoding 3D image information into a 2D light field image (LF), which is captured by a planar detection camera (DC). The sample 3D image is reconstructed from the LF image, based on the knowledge of the optical transformation. **(b)** SVIM improves LFM by illuminating a subset of the sample, through the use of light sheet (SPIM) illumination that is scanned axially, so that the thin sheet of light is extended into a slab. By illuminating only the relevant portions of the specimen, SVIM decreases background and increases contrast when compared to wide-field (WF) illumination of the entire sample. In the schematic presented here, the SVIM illumination axis is orthogonal to the detection axis, θ = 90^°^, but the benefits of reduced background can be obtained by using illumination from a different angle or by employing non-linear optical effects to selectively excite the volume of interest. **(c)** SPIM and SVIM 3D images of the trunk vasculature of a 5-dpf zebrafish embryo reveal different compromises between resolution and volumetric imaging time. SPIM offers a high-resolution 3-D rendering, but requires the collection of 100 sequential images and 99 movements of the collection plane; SVIM captures the same 3D volume in a single snapshot, at two orders of magnitude faster but with lower resolution. **(d)** SVIM contrast and effective resolution are improved as the illumination volume is decreased. Images of the same sub-volume of the GFP-labeled vasculature of a 5-dpf, *Tg(kdrl:GFP),* zebrafish larva show that SVIM with volume axial extent of 300 µm and 100 µm approach the performance of SPIM, and far exceed the image contrast obtained with wide-field illumination. The images shown are each average image projections of the same 40-µm thick sub-volume centered at approximately 170 µm into the specimen. **(e)** Quantitative comparison of image contrast, defined as the normalized standard deviation of the pixel value (Methods), comparing LFM, SPIM, and SVIM of different SVI extents from **(d)**. SVIM of increasingly smaller extents yields increasingly better contrast, quantitatively approaching the performance of SPIM. The contrast of SPIM shows the intrinsic contrast variation of the 3D sample, coupled with the expected decay for increasing depth, as the imaging plane gets deeper into the sample. The local increase in contrast seen for the SVIM cases at z = 0 µm comes from grid-like artefacts in the reconstructed images at the native focal plane. Supplementary Figures 3, 4, 5 perform more complete comparisons and analysis of these imaging datasets. Scale bars, 50 µm in **(c)**; 100 µm in **(d)**.

To better harvest the potential of LFM for the 4D analysis of biological tissues, we have developed an approach that limits the contribution from the specimen outside of the volume of interest. We combine LFM with selective illumination of the volume of interest (Fig. 1b), creating Light Field - Selective Volume Illumination Microscopy (SVIM). The simplest implementation of SVIM draws on the ability of Selective Plane Illumination Microscopy (SPIM; also known as light sheet microscopy) ^6^, which achieves low background and high contrast imaging by illuminating only the optical section of interest. SVIM expands the volume of excitation created by 1-photon or 2-photon SPIM to match the volume of interest by use of rapidly sweeping the light sheet (Fig. 1b) or by non-linear optics. By tailoring the directions of the illumination and detection, it is straightforward to minimize the amount of signal scattered into the volume of interest, thus reducing the background signal, increasing the contrast, and resulting in 3D reconstructions with greater overall image quality.

The implementation of SVIM tested here combined SVI and LFM modules with an existing custom-built SPIM ^7^. This permitted the same specimen to be imaged by both SPIM and SVIM so their capabilities could be directly and quantitatively compared (Methods, Supplementary Fig. 1). Our design of the light field detection module drew upon previous efforts ^3–5^, and the light field image reconstruction followed the 3D deconvolution approach described in Broxton *et al.* ^4^ using software made available by Prevedel *et al.* ^5^.

Imaging sub-diffraction beads over a ∼ 500 x 500 x 100 (x,y,z) μm^3^ volume showed that SVIM can capture 3D volumetric images in a single snapshot with ∼2 μm (FWHM) lateral resolution and ∼5 μm axial resolution (Supplementary Fig. 2), which is as expected for our experimental optical parameters (Methods). SPIM imaging of the same specimen, with the same primary magnification and numerical aperture, offered single plane images with ∼ 0.4 μm lateral and ∼ 1.5 μm axial resolution; thus, SVIM trades ∼ 3-5-fold in lateral and axial resolution for more than a 50-fold extension in z-depth coverage. We used SVIM to image in one snap shot the entire depth of the fluorescently-labeled trunk vasculature of the larval zebrafish, *Tg(kdrl:GFP)*, at 5 days-post-fertilization (dpf) (Fig. 1c). Compared to the 100-slice z-stack imaged with SPIM, the single-snap-shot SVIM captures faithfully the 3D structure of the GFP-labeled vasculature, with some softening of focus due to the expected reduced resolution. This demonstrates that with modest reductions in image quality, SVIM achieves dramatically increased z-depth coverage and imaging speed.

To study how the reduced volume of interest enhances image quality, we performed a series of SVIM acquisitions of the cranial vasculature of the same living 5-dpf zebrafish larva with differently-sized illuminated volumes of interest. SVIM image quality was characterized for illumination depth-extents ranging from 100 μm to the entire animal (with wide-field illumination) (Fig. 1d, Supplementary Fig. 3). SVIM image quality was far better than light field images from wide-field illumination, while falling short of the quality of SPIM imaging of the same specimen. As the z-extent of illumination decreased, background was reduced and contrast was increased. For quantitative image comparisons, we characterized the Image Contrast, defined as the standard deviation of the image pixel value divided by the average pixel value (Methods). The Image Contrast decreased as the z-extent of illumination increased, but it was higher for all SVIM conditions compared to wide-field illumination (Fig. 1e). The Image Contrast for the thinnest extent (100-μm) approached but fell short of the SPIM value.

The enhanced contrast of the thinner SVI volumes was observed throughout the axial extent of the captured volume (Fig. 1e). Importantly, the higher contrast of SVIM leads to better effective resolution, as demonstrated by the more faithful capture of the size of the imaged blood vessels (Supplementary Fig. 4). Finally, as the SVI sets the volume of interest, we confirmed that it is best to computationally reconstruct the illuminated volume in its z-entirety; if the volume is “under-reconstructed” with a smaller z-extent than its experimental SVI, poorer contrast is achieved (Supplementary Fig. 5). This is consistent with the underlying principle of the SVI approach, as any optical signal coming from the un-reconstructed z-planes will significantly increase the background term and degrade the final reconstructed image. This offers further validation of the benefit of SVIM for imaging in scattering biological tissues.

The synchronous volumetric imaging capability and the enhanced contrast of SVIM makes it an ideal tool to image dynamic biological systems where the components undergo fast motion in 3D space. We employed SVIM to perform 4D imaging of the bacterial flows in seawater, over a volume of 600 x 600 x 100 (depth) μm^3^ surrounding a Hawaiian bobtail squid, *Euprymna scolopes,* as its light organ is being selectively colonized by the bacteria *Vibrio fischeri* ^8,9^. The squid-bacteria symbiosis is an important model system for understanding the many facets of the interaction of bacteria and epithelial surfaces. The cilia-driven flow field around the light organ is critical for the initial capture of the bacteria by the ciliated surface of the light organ, leading to the eventual colonization of the host squid by the symbiont cells ^10^. However, previous measurements of the bacterial flow field were carried out in 2D ^11^, which inadequately captures the 3D flows around the light organ. Fig. 2a,b show the dramatically better image quality captured by SVIM as compared to conventional wide-field LFM. The excessive background of the wide-field LFM came from the extraneous excitation of the nearby tissues of the squid (Fig. 2a). SVIM removed enough of this background and enhanced the contrast to allow identification of the fluorescence from individual bacteria (Fig. 2b, Supplementary Movie 1). Analysis of the reconstructed SVIM 4D datasets yields the trajectories of individual bacteria, which permit a full and quantitative reconstruction of the flow-field in 3D (Fig. 2c, d, Supplementary Movie 2). The flow speeds reached up to 200 μm/s, far exceeding the *V. fischeri* self-propulsion speed of ∼ 70 μm/s ^12^. This is consistent with previous mutant studies showing that this initial stage of light organ colonization does not require endogenous motility of *V. fischeri.* ^13^, suggesting that the cilia-driven flow is likely the major driver of bacterial cell motion toward the light organ to initiate colonization. In future work we plan to deploy SVIM to quantify the 3D flow field as a function of various experimental conditions and perturbations, toward an understanding of both the conditions that set up the cilia-driven flow field, and how it contributes to the successful colonization of the squid host by the bacterial symbiont.

**Figure 2:**
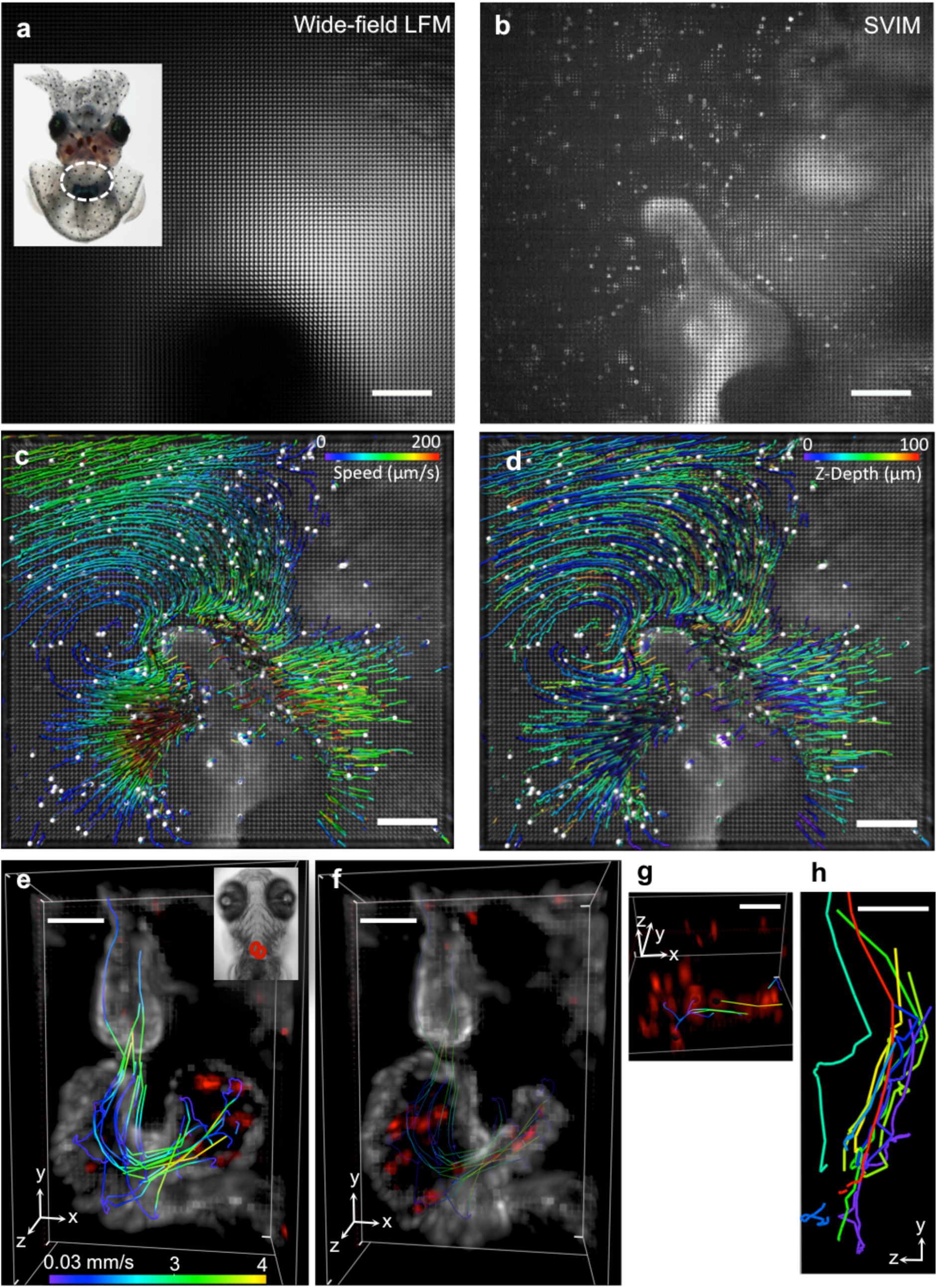
SVIM enables fast, high-contrast, volumetric imaging of live biological systems with fast 3D motion. We show examples in imaging the cilia-driven flow fields of *Vibrio fischeri* bacteria during their colonization of the light organ of the Hawaiian bobtail squid **(a-d)**; and imaging the blood flow in the beating heart of the zebrafish larva **(e-h)**. For both of these applications, the synchronous volumetric capability of SVIM is critical to capture the true 3D motion of the moving components. In the bacteria/squid application, **(a)** shows that raw light field images recorded with conventional wide-field illumination yielded excessive background, while **(b)** shows that SVIM, with the selective illumination volume of 100 µm, reduced this background and enhanced the contrast to allow localization of individual bacterium. Inset of **(a)** shows live juvenile squid, with region of light organ highlighted by dashed oval. **(c)** Quantitative flow trajectories tracked from the reconstructed data, color-coded for speed. **(d)** Quantitative flow trajectories tracked from the reconstructed data, color-coded for z-depth. The non-uniform 3D flow patterns were observed throughout the imaged volume, which would have been impossible to capture without SVIM. Images were collected at 20 volumes/s, of volume ∼ 600 x 600 x 100 (depth) µm^3^. (See also Supplementary Movies 1 and 2.) Scale bars, 100 µm **(a-d)**. In the application of imaging the wall motion of the beating heart and the moving blood cells **(e-h)**, a ∼ 250 x 150 x 150 (depth) µm^3^ volume of interest was imaged in a live 5-dpf zebrafish larva at 90 volumes/s; transgenes labeled the myocardium (rendered white) and blood cells (rendered red), *Tg(kdrl:eGFP, gata1:dsRed)*. Inset in **(e)** highlights the position of the heart. As shown in the volume renderings in **(e)** and **(f)**, the unprecedented high synchronous volumetric imaging rate and volume coverage enabled direct imaging and tracking of the quasi-chaotic blood flow, at single-blood-cell resolution throughout the cardiac beating cycle, which was about 450 ms. The moving endocardial wall was also faithfully captured, e.g. the atrio-ventricular (AV) canal and valve region were clearly identified. **(e)** At 133 ms after the AV valve opened there are rapid blood-cell movements. Representative blood-cell trajectories through the heart were manually tracked and quantified (color of the trajectory depicts blood-cell speed). **(f)** At 244 ms after the AV valve opened the heart has a dramatically different shape and blood flow. **(g)** Perspective view of the blood cells demonstrates the achieved single-cell resolution, notably along the z direction. **(h)** Maximum projection image along the x-axis of some representative flow trajectories highlights the significant component of blood flow along the z direction. To aid visualization, clipping planes in the y-z plane have been used to cut out the atrium and parts of the ventricle. Color-coding of the blood-cell tracks here is for visual identification. (See also Supplementary Movies 3 and 4.) Scale bars, 50 µm **(e-h)**.

A further test of SVIM is provided by imaging the motions of the live beating heart of zebrafish larvae at 5 dpf (Fig. 2e-h, Supplementary Movies 3 and 4), in the effort to understand the roles that dynamic cellular and fluid motions contribute to the vertebrate heart development ^14,15^. Methods based on retrospective synchronization ^7,16,17^ and prospective gating ^18^, all based on the assumption that the heart beating is periodic, have allowed the reconstruction of heart wall motions. These approaches, however, cannot capture the quasi-chaotic 3D flow field of the blood. SVIM, with its unique synchronous volumetric imaging capability, permits direct imaging of blood cells as they travel through the entire 3D heart (Fig. 2e, f), resolving individual blood cells even during the window when they travel at the fastest speed of up to 4 mm/s through the atrioventricular canal (Fig. 2g). The high 4D spatiotemporal resolution of our SVIM allows us to track blood cells’ flow trajectories with single-cell resolution (Fig. 2e), showing a distinct quasi-chaotic nature of the flow field, with significant motion component in the z-depth direction (Fig. 2h), which would have been missed without the extended depth-coverage of SVIM. Further, the imaging speed, coverage, and resolution demonstrated here, 90 volumes/s with volume ∼ 250 x 250 x 150 (depth) μm^3^ at single-blood-cell resolution, is impossible to achieve with the sequential fast plane-scanning approach ^19^, as an imaging speed > 5000 frames/s would be required. This high imaging speed is not only hardware-limited but also would induce excessive photo-damage due to the intrinsic low duty cycle of the sequential scanning approach. In capturing the wall motion of the beating heart, the reduced resolution of SVIM, Fig. 2e-f, does not yield the sub-cellular resolution achieved with retrospective synchronization methods ^7,17^. However, the synchronous volumetric nature of SVIM means that it captures faithfully both the motion and changing signal intensity of the dynamic tissue along the imaging axial (z) axis, free from potential artifacts caused by the assumption of signal continuity along the axial axis of current retrospective synchronization methods ^7,16,17^. Thus, with the capability of directly recording the unbiased 3D motion of the beating heart and the result blood flow, we envision SVIM will be an important tool in characterizing the dynamic properties of the developing heart, potentially extending recent works that map electrical signaling over the heart ^20^ or link cardiac valve formation to blood flow dynamics ^21–23^.

Functional neuro imaging is another area where the high-contrast, synchronous volumetric imaging capability of SVIM could provide game-changing benefits. We used SVIM to image the cellular brain activity of zebrafish larvae, labeled with pan-neuronal fluorescent calcium indicators (Fig. 3). The higher contrast of SVIM, compared to wide-field illumination, enables cleaner capture of single-neuron firings throughout the imaged volume (Fig. 3a). Furthermore, we implemented SVIM with 2-photon (2p) excitation in a straightforward manner, akin to 2p-SPIM ^24^, by fast scanning in 2D the extended excitation focus to cover the entire 3D volume of interest at least once during the exposure time of the camera (Methods, Supplementary Fig. 1). Better contrast is achieved with 2p-SVIM than 1p-SVIM, Fig. 3a, due to not only the benefits of nonlinear excitation ^24^, but also because the near infrared light (NIR) used in 2p-SVIM does not illicit a direct visual response in the zebrafish as the visible light used in 1p-SVIM does ^25^, leading to less background calcium response in the brain when 2p-SVIM is used.

**Figure 3:**
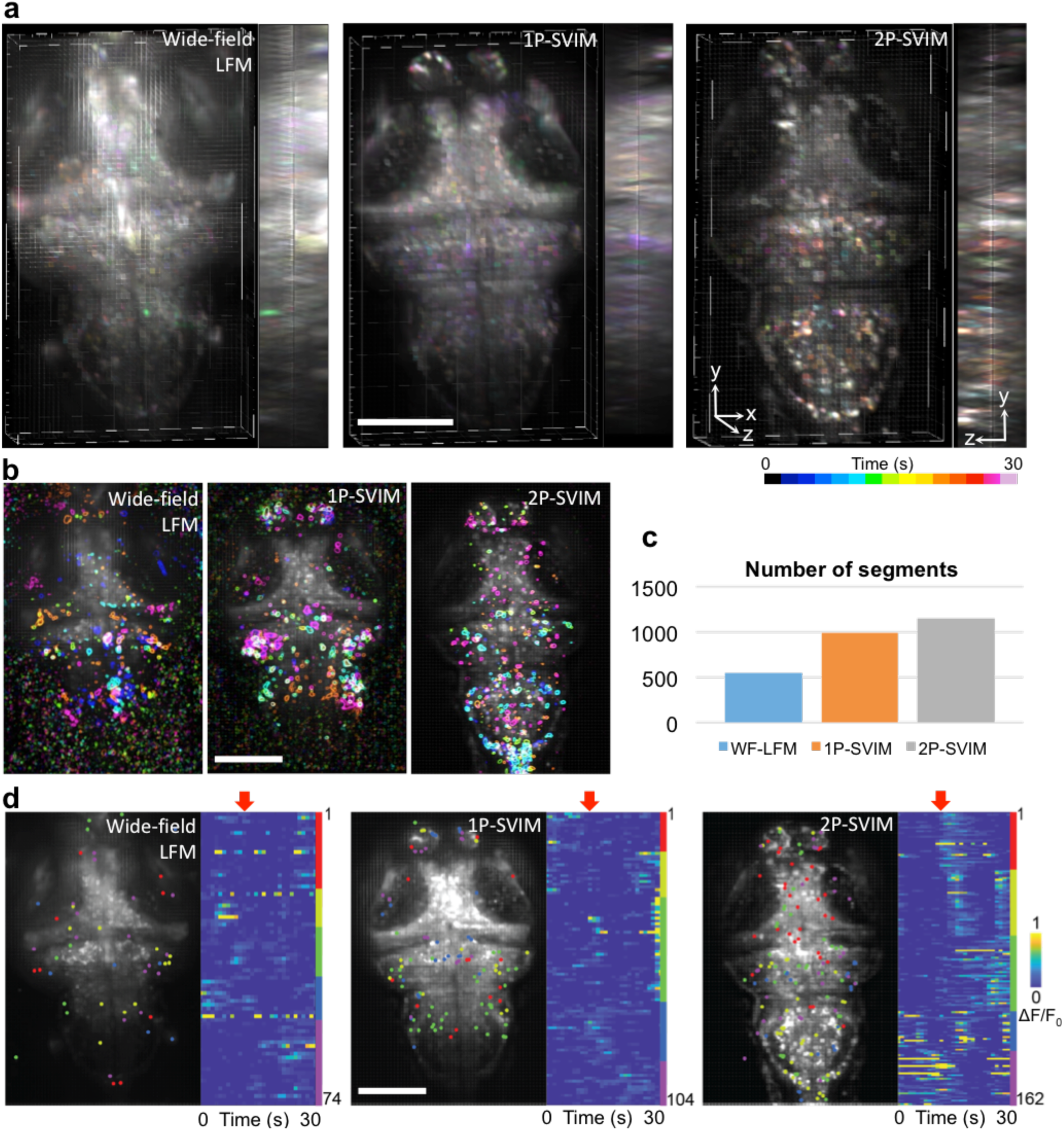
Enhanced contrast of SVIM enables capture of more active neurons in brain-wide functional imaging. **(a)** SVIM, in either 1-photon-excitation (1p-SVIM) or 2-photon-excitation (2p-SVIM) modes, produces superior contrast to the conventional 1-photon wide-field LFM, in functional imaging of 7-dpf larval zebrafish with pan-neuronal fluorescent calcium indicators, *Tg(elavl3:H2b-GCaMP6s)*. Images shown are 3D-rendered views of the reconstructed datasets, with temporal color-coding of the signal intensity over a 30-s time window of spontaneous brain activity, highlighting the single-cell resolution achieved with the SVIM approaches. **(b)** Principal component analysis (PCA) and independent component analysis (ICA) of reconstructed images yielded regions of brains that exhibit independent temporally varying calcium transients. Shown are five representative ICs for each of the imaging modality. The best high quality, cell-like ICs, with less noise, were obtained by 2p-SPIM, followed by 1p-SVIM, and then wide-field LFM. **(c)** Segments, consisting of independent components spatially filtered through the expected size of single neurons, represent computationally extracted single, active neurons. The enhanced contrast of the SVIM approaches over wide-field illumination yielded approximately two times more extracted segments, with the most segments achieved via 2p-SPIM due to its highest contrast. **(d)** Light-evoked activity extracted using PCA/ICA, where each dot represents an extracted single-neuron segment. K-means clustering separates the activity traces of segments into functionally distinct groups that respond differently to light onset (red arrow). Only the dataset collected with 2p-SVIM exhibited a clear response to the visual stimulus, due to the invisibility of the 2p excitation light (see text for details). Scale bars, 100 µm.

We carried out Principal Component Analysis and Independent Component Analysis (PCA & ICA) ^26^(Methods) to computationally extract single-cell responses from the 3D reconstructed datasets and provide a quantitative measure of the improved performance of SVIM over wide-field illumination. Single-neuron responses were successfully extracted from all modes of imaging for a 30-sec window of spontaneous brain activity (Fig. 3b), but, importantly, the higher contrast of the SVIM approach allows a factor of two more single-neuron segments to be extracted compared with the wide-field case (Fig. 3c). We also examined the brain activity of the zebrafish larva as it was subjected to the effect of a light source being turned on (Methods) (Fig. 3d). Application of k-means clustering to the temporal response of the extracted segments reveal correlated activity of a variety of neuronal clusters, but only the brain activity imaged with 2p-SVIM reveals a distinct light-evoked response, demonstrating the advantage of 2p illumination in avoiding interference with the visual response of the animal, as discussed above, due to the invisibility of NIR light.

In summary, we have demonstrated that, by combining the strategy of selective volume illumination to light field microscopy, SVIM provides a unique tool for high-contrast synchronous volumetric imaging of dynamic systems. Many innovations in the last decade have revolutionized 4D imaging of living tissues, employing fast sequential scanning of the imaging optical plane to capture cellular details across the sub-micron to mesoscopic scale ^27–32^. SVIM expands this 4D imaging toolkit in a unique direction, with its capability of capturing an extended 3D volume in a single snapshot with high contrast, with modest sacrifice in resolution. The applications of imaging the bacterial-flow around the squid light organ and blood flow in the beating heart, Fig. 2, demonstrate SVIM to be the enabling imaging technology in imaging and tracking 3D flows and moving tissues, where it is critical to capture the 3D image synchronously to avoid any bias in the derived 3D dynamics. As LFM has found more applications in neuro imaging in recent years ^5,33–38^, the demonstrated higher contrast of SVIM compared with wide-field LFM in functional imaging of neurons, Fig. 3, should enable an overall improvement in the information content captured. The fast volumetric imaging speed of SVIM should make it a particularly attractive tool for neuro imaging with the next generation of voltage-sensitive sensors ^39,40^, and for functional imaging of behaving animals, taking advantage of preparations and imaging platforms where the sample is unrestrained ^33,38,41–43^.

We have implemented SVIM with simple geometrical manipulation of the illumination and detection axes, and with nonlinear optical excitation. Compared to SPIM, SVIM requires less stringent optical requirements – as long as a significant portion of extraneous illumination is removed, the SVIM benefit will ensue. SVIM improves on conventional LFM by optimizing the illumination pathway, and thus is fully compatible with recent innovations in the detection pathway of LFM, via manipulation the phase content of the detected light field ^44,45^ or capturing the light field at the image plane ^33^. LFM belongs to an emerging class of computational imaging techniques that utilize the power of physical modeling, signal processing, and computation to enable new performance space beyond conventional microscopy ^34,37,46–48^. Since the computational speed and robustness will benefit from higher contrast in the raw imaging data, we envision that the general strategy of contrast enhancement by selective volume illumination will play a key role in bringing the benefits of computational imaging to a wide variety of biomedical applications.

## Acknowledgement

We thank EG Ruby (University of Hawaii-Manoa) for critical assistance and sharing of resources for the squid-bacteria experiments; Misha Ahrens (Janelia) for sharing of zebrafish lines.

Funding was provided for by the Gordon and Betty Moore Foundation grant #3396 (Ruby, McFall-Ngai and Fraser); National Institute of Health, TR01 grant #1R01MH107238-01 (Arnold, Kesselman, Fraser), R01 grant #AI150661 (McFall-Ngai and Ruby), grant #OD11024 (Ruby and McFall-Ngai); NSF BRAIN grant #1650406 (Dickman, Fraser, Truong); NSF INSPIRE grant #1608744 (Kanso, Fraser). SM was supported by the USC Provost Fellowship; KKD by the Alfred Mann Institute Doctoral Fellowship.

## Methods and Supplementary Materials

### Methods 1 - Microscopy setup and implementation

The optical setup was based on an existing SPIM setup^7^, with modifications to provide the selective volume illumination and light field detection (Supplementary Fig. 1, Supplementary Table 2). Briefly, 1p-excitation continuous-wave lasers and 2p-excitation femtosecond-pulsed laser light were combined and directed at the sample through a pair of galvanometer scanners and scanning optics. The fluorescence signal was collected in the direction orthogonal to the illumination axis, through appropriate spectral optical filters, and detected with a detection module that allowed imaging either in SPIM or SVIM mode. For SVIM mode, a micro-lens array was placed at the conventional image plane to capture the light field coming from the sample, which was subsequently recorded by the detection camera^3,4^. Computer-controlled motorized stages were used to allow reproducible switching between SPIM and SVIM modes. To provide selective volume illumination, the galvos controlling the illumination light were adjusted to paint out the desired illuminated volume within a single camera exposure. Image acquisition was through the software Micro-Manager^49^ and a custom software written in Labview (National Instruments). See Supplementary Fig. 1 and its caption for more detailed descriptions and Supplementary Table 1 for part numbers of key components.

### Methods 2 – Sampling handling and imaging procedure

All animal handling procedures follow guidelines established in the Guide for the Care and Use of Laboratory Animals at the University of Southern California. All zebrafish lines used are available from ZIRC (zebrafish.org).

#### Zebrafish experiments

zebrafish embryos were collected from mating of appropriate adult fish and raised in egg water (60 µg/ml of stock salts in distilled water) at 28.5 °C. At 20 hpf, 1-phenyl-2-thiourea (PTU) (30 mg/L) was added to the egg water to reduce pigmentation in the animals. For imaging experiments, the samples were embedded in a 1 mm-diameter cylinder of 1.5% low-melting agarose (SeaPlague) for imaging in the SPIM/SVIM setup, as described in^7^. The imaging chamber was filled with 30% Danieau buffer (1740 mM NaCl, 21 mM KCl, 12 mM MgSO_4_•7H_2_O, 18 mM Ca(NO_3_)_2_, 150 mM HEPES) at 28**°**C. Anesthetics was used (Tricaine, 100 mg/L) during the mounting procedure and imaging, except for during neural functional imaging. For structural imaging of the cranial vasculature, 5-dpf larvae, *Tg(kdrl:eGFP)*, were used. For heart imaging, transgenic adults, *Tg(kdrl:eGFP)* and *Tg(gata1:dsRed)*, were crossed to produced offsprings with fluorescent labels in both the blood cells (dsRed) and endocardium (eGFP). Samples, at 5 dpf, were selected for heterozygous expression in gata1:dsRed to reduce the density of fluorescent blood cells. Two-color imaging of the blood cells and endocardium were carried out sequentially, at rate of 90 volumes/s. For neural functional imaging, zebrafish larvae at 7 dpf with nuclear-localized pan-neuro fluorescent calcium indicators, *Tg(elavl3:H2B-GCaMP6s)*, were imaged at rate of 1 volume/s. The recording was for 300 s, with evoking-light turned on during t = 99 to 199 s, provided by a 625 nm LED (Thorlabs).

#### Squid/bacteria experiments

squid samples were procured as previously described^50^. Briefly, adult squids were collected in Oahu, Hawaii, and then housed and allowed to reproduce at a facility at the University of Wisconsin, Madison. Hatchling squids were shipped overnight to our laboratory, and used for experiments within 3 days of arrival. A solution of 2% ethanol/artificial seawater (Instant Ocean) was used to anesthetize the squids during mounting and imaging. Under a stereo dissection microscope, the mantle of the squid was carefully cut open and trimmed to expose the light organ of the animal. Then the animal was embedded in a cylinder of agarose (2% low melt agarose, 2% ethanol, in artificial seawater) using a procedure similar to that used for zebrafish. The cylinder and plunger unit were fashioned from a FEP tube (inside diameter ∼ 2mm) and glass rod (outside diameter ∼ 2mm), respectively, to accommodate the size of the squid. To allow direct interaction between the light organ and the surrounding fluid environment, the small volume of solidified agarose encasing the light organ was carefully removed with forceps. The agarose-embedded squid was then mounted into the imaging chamber (containing 60 mL of the ethanol/seawater solution), and GFP-expressing *V. fischeri* ES114 carrying pVSV102^51^ was added to reach a concentration of 50,000 cfu/ml. The squid and bacteria were monitored using SPIM for approximately 2 hours before imaging of the bacterial flow fields with SVIM was carried out.

### Methods 3 – Light field image processing and 3D reconstruction

Reconstruction of light field images was carried out using the wave-optics procedure described in^4,5^ and the software package made available by^5^, and with the ray-optics software LFDisplay^3^. Briefly, the rectifying parameters of the acquired light field images, describing the geometrical relationships between the micro-lens array and the detection optical train, were found in LFDisplay. Theoretical point spread functions (PSF) were calculated using the optical parameters of the entire imaging path, and the desired spatial sampling and coverage of the 3D reconstruction^5^. Then, the rectifying parameters and the PSF were used as inputs into the 3D wave-optics reconstruction program from^5^ to reconstruct the acquired 2D light field images into 3D images. Two key parameters for the PSF calculation and the resulting 3D reconstruction are the z-extent of the volume to be reconstructed and the desired z-sampling (i.e. thickness of individual z-slice). Larger z-extent and finer z-sampling requires more onboard memory for the graphical processing unit (GPU) used in the 3D reconstruction program. With the GPU that we used (Titan X, Nvidia), the largest z-extent that we could reconstruct was 400 µm, at z-sampling of 2 µm. Consequently, for results shown in Fig. 1d, e and Supplementary Figs. 3 and 4, both datasets of 550-µm SVIM and WF-LFM were reconstructed with z-extent of 400 µm, while all others were reconstructed with z-extent equal to their actual experimental illumination extent. Similarly, 400-µm was the largest reconstruction extent that we could carry out for the results shown in Supplementary Fig. 5.

We used the same micro-lens array (pitch = 150 µm, focal length = 3 mm) for two primary imaging conditions of (32X magnification, 0.8 NA) or (20X magnification, 0.5 NA). With these parameters, and the reconstruction parameters listed in Supplementary Table 1, following^4^ we expect to have the following approximate resolution for our reconstructed datasets: for the 32X magnification case, 2 (& 5) µm laterally (& axially); for the 20X magnification case, 3.5 (& 12) µm laterally (& axially).

### Methods 4 – Image analysis and presentation

SPIM images were background-subtracted to account for camera dark-counts. SVIM images were background-adjusted within the reconstruction process. Then all SPIM and SVIM images were scaled to fill the full 16-bit dynamic range. For visualization in the figures, image pixel intensities were further scaled to minimum and maximum display contrast with 0.4% saturation. Unless otherwise noted, 2D image processing and analysis were done in Fiji^52^, while 3D rendering and analysis were done in Imaris (Bitplane).

#### Zebrafish vasculature

For zebrafish vasculature images (Fig. 1d, Supplementary Figs. 3 and 5), the 3D datasets were displayed as an averaged projection in z, for the same volume section extending from z = - 48 to ‒12 µm, where z = 0 µm is the native focal plane of the detection objective. This volume excludes the native focal plane of the imaged z-stack, where grid-like artifacts from the light field reconstruction are most prominent. The native focal plane was experimentally set at approximately 200 (or 150) µm into the zebrafish head from its dorsal surface, for datasets shown in Fig. 1d and Supplementary Fig. 3 (or Supplementary Fig. 5).

#### Bacteria/squid

For squid/bacterial results, Fig. 2a-d, Supplementary Movies 1 and 2, tracking and quantification of the bacterial flow field were carried out using the automatic spot segmentation and tracking functions in Imaris, followed by manual correction.

#### Zebrafish heart/blood

For results describing the zebrafish beating heart, Fig. 2e-h, Supplementary Movies 3 and 4, the sequentially-acquired light field time-series data of the endocardium and blood flow were reconstructed separately. The reconstructed 4D datasets, each spanning approximately 4 heart beats, were then synchronized in time by renumbering the endocardium frames such that the time point at which the atrium is most contracted in the endocardium movie matches the time point when, in the blood flow movie, the flow into the ventricle from the atrium momentarily stops. After synchronization, the two movies were overlaid to create a composite 2-color movie. Tracking and quantification of the blood cells’ flow trajectories in the zebrafish heart were carried out manually in Imaris. Twelve representative blood cells were tracked and quantified.

#### Zebrafish brain activity

For the zebrafish brain activity results, Fig. 3, the analysis was carried out using a combination of Fiji, Imaris, and MATLAB. In Fig. 3a, to represent temporal data, we performed temporal color-coding using the “spectrum” color-map in Fiji over a 30-second time window of spontaneous brain activity, and the resulting color-coded images were rendered in Imaris for visualization. In Figs. 3b and Fig. 3d, reconstructed data were displayed as average intensity projection, overlaid with representative extracted independent components and segmented cells, respectively.

Principle component analysis (PCA) and independent component analysis (ICA) of the brain activity datasets, acquired by different imaging modalities, was carried out using the MATLAB-based framework from^26^. Briefly, 30-s time windows of time series data were first dimensionally reduced by PCA. We found that 90% of the dataset variance was covered by 20 principal components (PC), for all imaging modalities. ICA was then applied to these 20 PCs to find 20 independent components (IC), which denote spatial domains that exhibit independent time-varying behavior. Five representative ICs were shown for each of the imaging modality in Fig. 3b. To find spatial segments that represent individual active neurons, we smoothed the IC results with a Gaussian blur of 6-µm diameter, applied a threshold of 2.3 standard deviations, and then working with yz images we applied an elliptical segmentation mask of 5 µm x 12 µm (minor x major axes, along y and z directions, respectively) to find the segments as reported in Fig. 3c. The elliptical mask corresponded to the expected size of individual neuronal cell nucleus (∼ 5 µm diameter) convolved with the resolution achieved with the SVIM imaging and reconstruction parameters (∼ 3.5 and 12 µm, along y and z, respectively).

The same PCA/ICA framework was used to analyze the neural activity signals corresponding to the light-evoked response, Fig. 3d. To find approximated single-cell activity, segmentation was done on IC filters on maximum intensity projection images in xy, and the found segments, represented as colored dots in Fig. 3d, were used to extract temporal activity traces. K-means clustering and linear correlation of the segments’ activity traces were then carried out in MATLAB. Using the Calinski-Harabasz clustering evaluation criterion, we found that the optimal K was 5 for datasets from all imaging modalities. Neural activity traces were arranged according to clustering result. Heat map panels in Fig. 3d showed the normalized fluorescent activity traces from segmented cells, presented as ΔF/F_0_.

### Methods 5 – Comparative analysis of image contrast and effective resolution

#### Contrast

To compare image contrast between SPIM, SVIM, and wide-field LFM, we measured the relative standard deviation of the pixel intensities from the respective images. The standard deviation *σ* of an image, which is the same quantity as root-mean-square (RMS) contrast that appears in vision science^53,54^, is given by

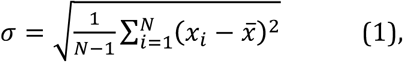

in which *x*_*i*_ is a normalized intensity-level value, so that 0 ≤ *x*_*i*_ ≤ 65,535, and *x* is the average intensity of all pixel values *N* in the image:

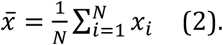

Putting the expressions above together, we have

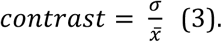

We emphasize that this measure of image contrast is independent of the total pixel count *N*, and thus provides a concise metric for direct comparison of contrast between SPIM, SVIM with various SVI extents, and wide-field LFM, at any z-position in the volume of interest (Fig. 1e and Supplementary Fig. 5f). Analysis was performed in Matlab.

#### Effective resolution

To evaluate the effective resolution between SPIM, SVIM with various SVI extents, and wide-field LFM, the intensity profiles of representative blood vessels were extracted from a single plane in the imaged volume for all modalities (Supplementary Fig. 4). The line segments were generated using a Matlab script, which specified paths perpendicular to the direction of the blood vessels. Peak values along the line segments thus correspond to the brightest blood vessel structures. In all the intensity profile plots, the intensity values were normalized and fitted by the global maximum. All raw and processed data, and custom Labview and MATLAB scripts/codes, are available upon request from the authors.

---

**Supplementary Figure 1.**
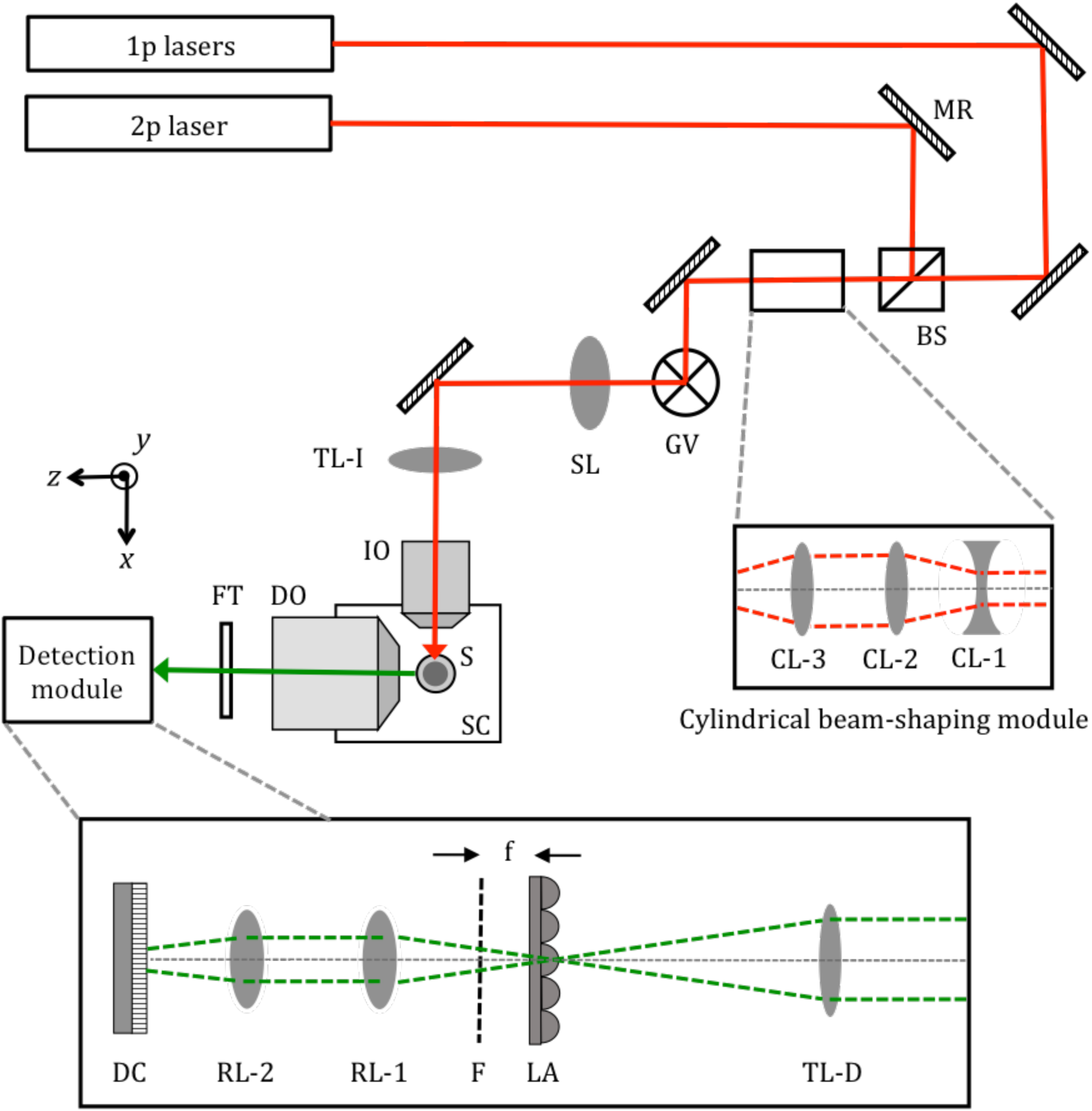
Schematic of the SVIM/SPIM optical setup, and descriptions of operation procedure. Top-view of the optical setup, depicting only the key components. (See Supplementary Table 1 for specific part numbers of the key components.) Laser light, for 1-photon (1p) and 2-photon (2p) excitation, were directed by silver mirrors (MR) and combined into the same optical path by a polarizing beamsplitter (BS). The laser light was then routed into a 2-dimensional scanning galvos module (GV), then through scan lens (SL) and illumination tube lens (TL-I), and into illumination objective (IO). Sample (S) was suspended from the top of the liquid-filled sample chamber (SC), and detection objective (DO) collected the fluorescence light generated at the sample. The y direction of the experimental coordinate system is anti-parallel to gravity. The beam at GV was imaged to the back focal plane of IO by the SL and TL-I, so adjusting the rotational positions of the galvos enabled translating the illumination beam at the sample in the y and z directions. Fluorescence signal collected by the DO passed through the appropriate interference bandpass filters (FT) that blocked the laser light and selected for the right detection colors. The fluorescence was then recorded by a detection module that allowed imaging in either SVIM or SPIM mode. In SVIM mode, as depicted in the figure, detection tube lens (TL-D) formed the image of the sample at its focal plane, where micro-lens array (LA) was placed. The desired light field image was then formed at focal plane F, at distance f away from the LA (where f = focal length of the micro-lens). A pair of identical photographic lenses (RL-1 and RL-2), configured in 4f mode, relayed the light field image at F to be recorded at detection camera (DC). The optical parameters of the LA and the DO and TL-D (see Supplementary Table 2) were chosen to ensure that the spatial-angular bandwidth of the light field collected by the DO was matched to that of the LA (Levoy 2006). To operate in SPIM mode, LA was moved entirely out of the optical path, and the entire block of (RL-1, RL-2, and DC) was moved by distance f closer to the TL-D (along the –z direction). Computer-controlled motorized translational stages (not shown) were used to allow convenient and reproducible switching between SVIM and SPIM modes. During SPIM imaging, the illumination beam was scanned in the y direction, resulting in a scanned light sheet transecting the sample in the xy plane, and the sample was moved along z to achieve 3D imaging. During SVIM imaging with 2p excitation, the beam was scanned in both the y and z directions, with appropriate amplitudes, to “paint” out the desired selective volume illumination (SVI) at the sample. During SVIM imaging with 1p excitation, a removable cylindrical beam-shaping module was put in place, up-stream of the GV, where cylindrical lenses (CL-1) and (CL-2) expanded the beam along y-direction by 2x, and cylindrical lens (CL-3) focused the beam along the y-direction at the GV. This focusing resulted in a static illumination light sheet (in the xy plane) at the sample, and the beam only needed to be scanned along one direction, z, to enable painting out the desired SVI. A custom-written Labview program, in conjunction with the software Micro-Manager (Edelstein 2014), coordinated the galvos scanning and camera triggering to ensure that within one camera exposure the volume of interest was illuminated an integer number of times, or that it was illuminated > 10 times, to ensure that the excitation intensity was uniform to better than 10% from frame to frame. Not shown was an imaging module looking at the sample from opposite to the IO, which allowed observation of the laser illuminated sample region in the yz plane. This imaging module was used to verify and calibrate the galvos scanning parameters to achieve the desired SVI. Also not shown was a separate beam path that directed the illumination laser light toward the sample along the +z direction – this path was used for wide-field illumination of the sample.

**Supplementary Figure 2.**
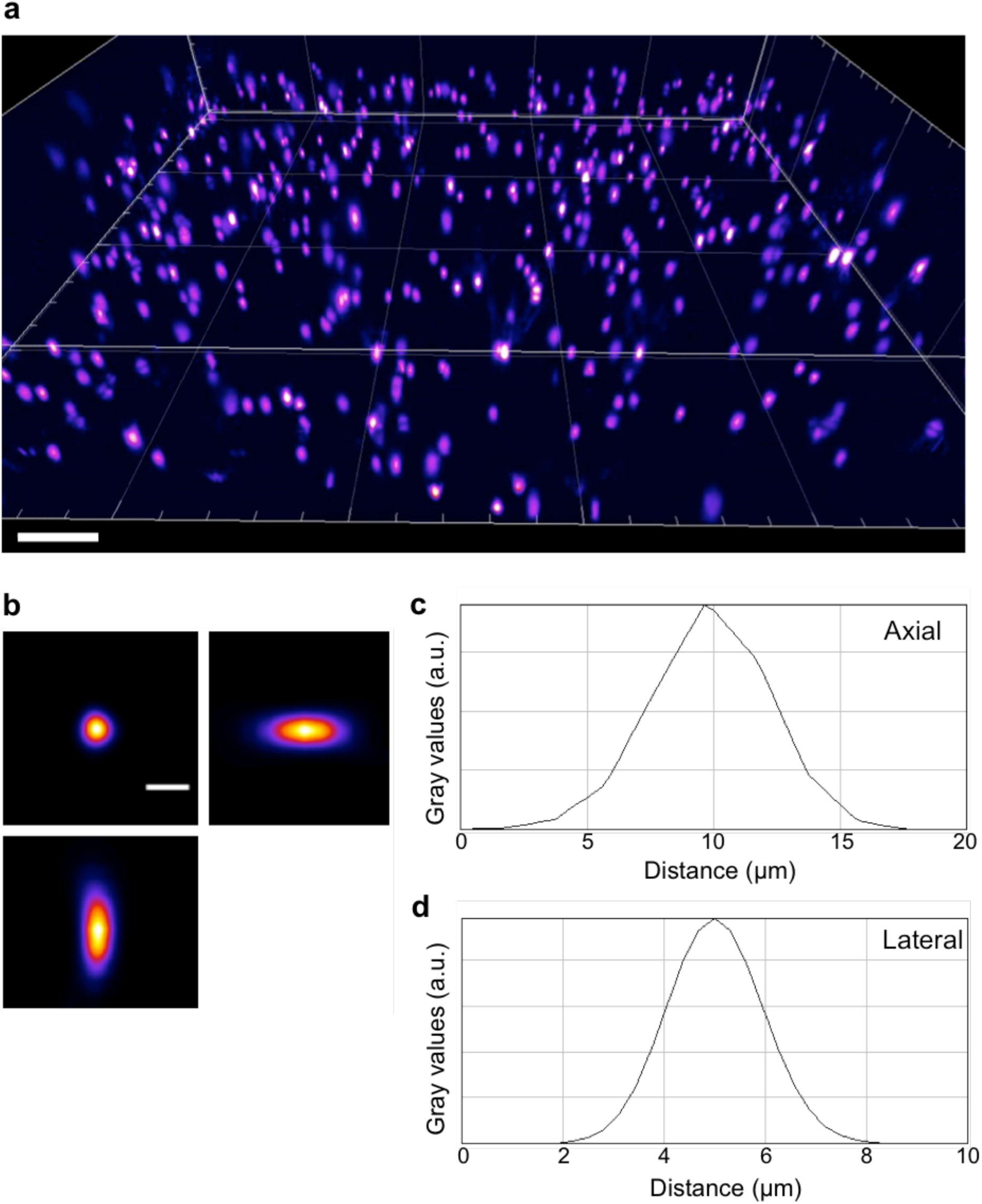
Imaging of sub-diffractive fluorescent beads to characterize resolution of SVIM. Resolution performance of SVIM was characterized by imaging sub-diffractive fluorescent beads (diameter = 175 nm, PS-Speck, ThermoFisher) embedded in low-melting agarose. **(a)** 3D-rendered view of the reconstructed volume of beads, spanning ∼ 500 x 500 x 100 (x,y,z) μm^3^. Ortho-sliced images of a representative bead image are shown in **(b)**. Line profiles along the lateral **(c)** and axial **(d)** directions show an approximate FWHM resolution of 2 μm laterally and 5 μm axially. This was as expected for the optical parameters used (see Methods). Scale bars, 50 μm **(a)**, 5 μm **(b)**.

**Supplementary Figure 3.**
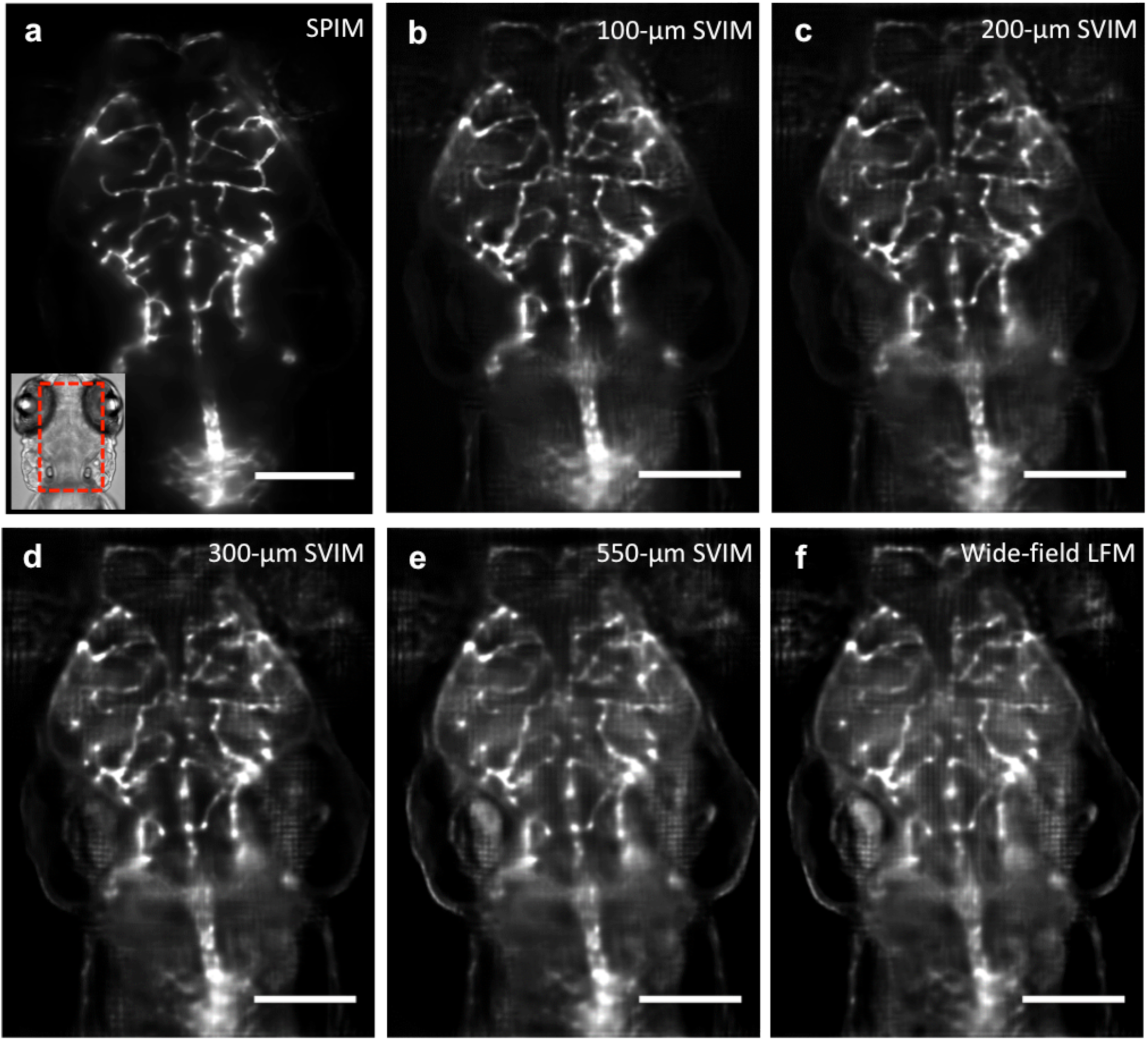
SVIM offers improved contrast and effective resolution over conventional wide-field LFM. (This figure is the expanded version of Fig. 1d.) Images of the same sub-volume of the GFP-labeled vasculature of a 5-dpf, *Tg(kdrl:eGFP),* zebrafish larva captured with **(a)** SPIM, and **(b-f)** SVIM with z-extents of 100, 200, 300, and 550-µm, respectively, and **(g)** wide-field LFM. SVIM of increasing smaller z-extents have increasingly higher contrast and effective resolution, approaching the performance of SPIM, and far exceeding the image quality obtained with wide-field illumination. The images shown are each average image projections of the same 40-µm thick sub-volume, centered at approximately 170 µm into the specimen. Scale bar, 100 µm.

**Supplementary Figure 4.**
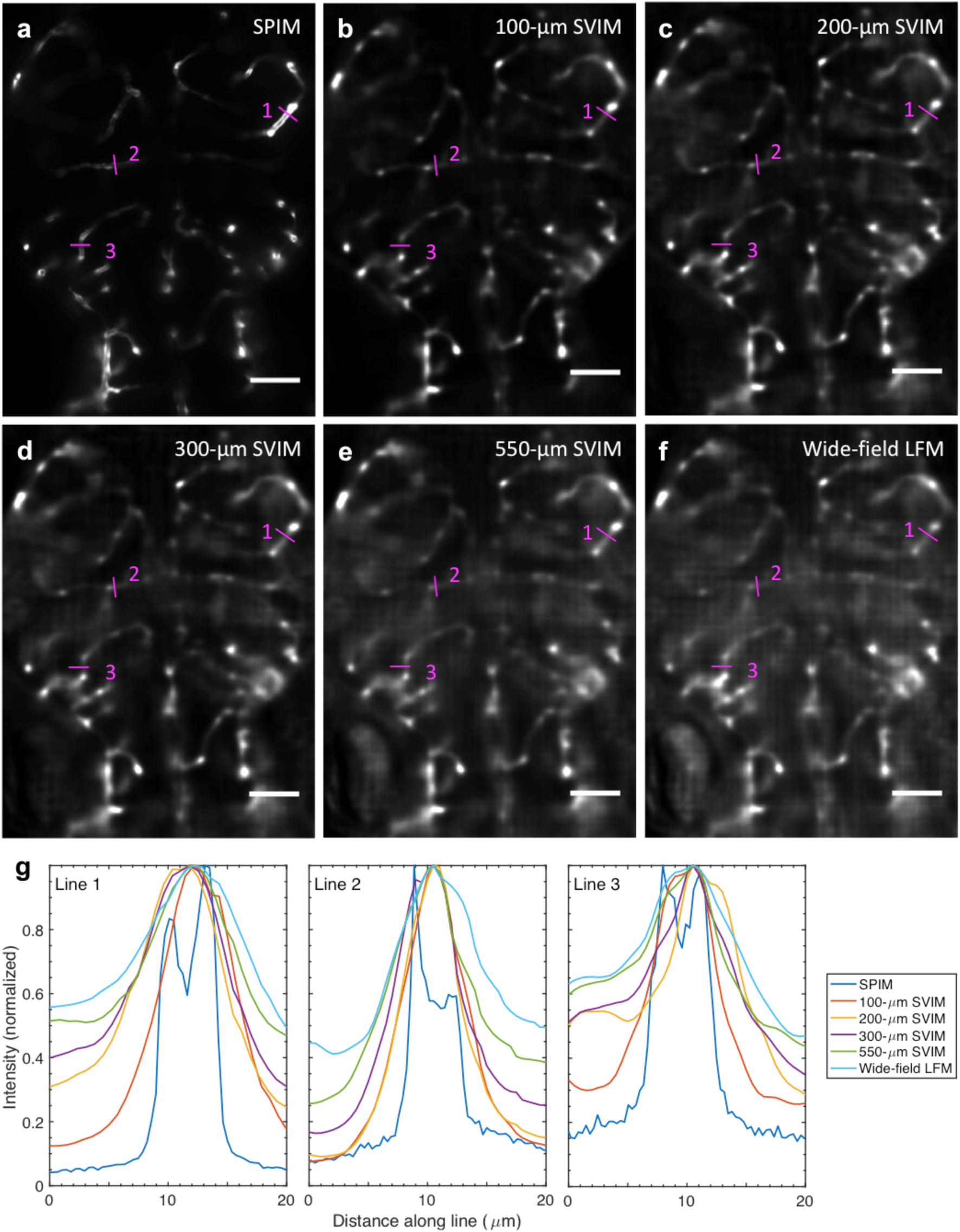
Enhanced effective resolution enabled by the better contrast achieved by selective volume illumination. Single-plane (relative z = ‒48 μm) images of the trunk vasculature of a 5-dpf zebrafish embryo, taken from **(a)** SPIM, **(b-e)** SVIM, and **(f)** wide-field LFM. **(g)** Intensity plots along the three magenta lines which traverse individual blood vessels, as indicated in the images. In all three plots, the intensity values were normalized and fitted by the global maximum. SPIM, as expected, exhibits the best resolution, resolving the separate sidewalls of the vessels. With SVIM, smaller SVI extents yield narrower peaks, decreased background, and increasingly better resolution, approaching the ground-truth established by SPIM. Scale bars, 50 μm.

**Supplementary Figure 5.**
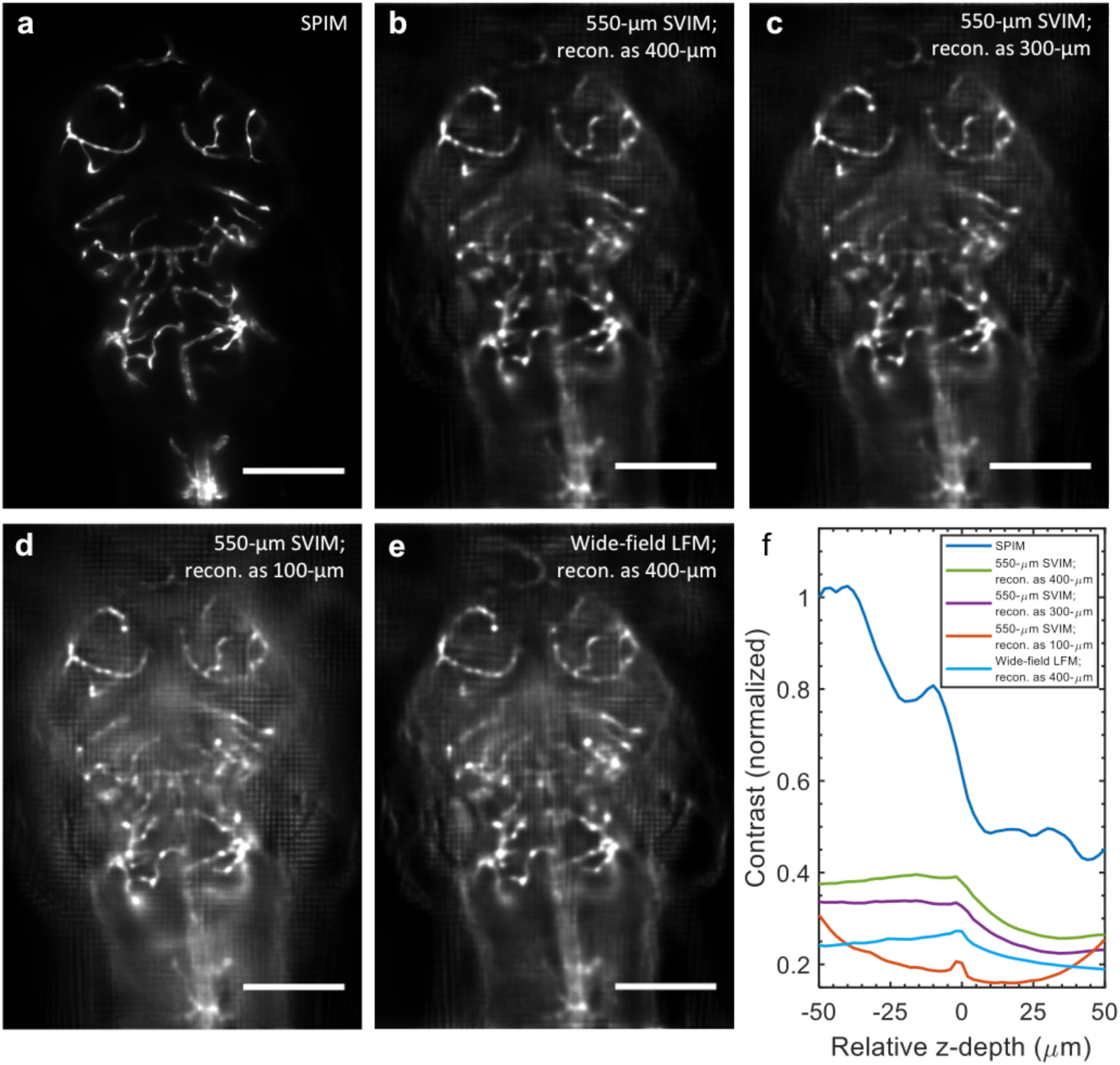
SVIM contrast is degraded when images are “under-reconstructed” as volumes smaller than the illuminated volume. Images of the same 40 µm-thick sub-volume of the GFP-labeled vasculature of a 5-dpf, *Tg(kdrl:eGFP),* zebrafish larva, shown as average image projections, centered at approximately 120 μm, comparing **(a)** SPIM, **(b-d)** 550-μm SVIM reconstructed as volumes of various z-extents (400 μm, 300 μm, and 100 μm), and **(e)** wide-field LFM reconstructed as 400-μm z-extent volume. For each modality, **(f)** shows the quantitative image contrast measured as function of z depth and normalized against the top plane (z = ‒50 μm) of SPIM. A moving average was applied to compute the SPIM contrast curve, as in Fig. 1e. As expected, SVIM with under-reconstructed (300-μm and 100-μm) z-extents have poorer contrast than SVIM reconstructed as volumes approaching its entire illuminated volume (400-μm). Further, 550-μm SVIM under-reconstructed with a 100-μm z-extent exhibits even lower contrast than wide-field LFM. (Note that the up-tick in contrast for the case of (550-μm SVIM reconstructed as 100-μm z-extent) toward the edges of the volume comes from high-contrast reconstruction artifacts). Scale bars, 150 μm.

**Supplementary Table 1.** Part numbers and descriptions of key components of optical setup.

Part numbers and descriptions of key components of optical setup.
| Item | Acronym<br>(Suppl. Fig. 1) | Detailed descriptions | Part number | Manufacturer/<br>Vendor |
| --- | --- | --- | --- | --- |
| Laser system, visible 1-photon excitation | 1p lasers | SOLE-6 Light Engine (lines used: 488, 561 nm) | SOLE-6 Light Engine | Omicron |
| Laser, near-infrared 2-photon excitation | 2p laser | Chameleon Ultra 2 | Chameleon Ultra 2 | Coherent |
| Silver mirrors | MR |  | PF10-03-P01 | Thorlabs |
| Polarizing beamsplitter | BS | Broadband, 620-1000 nm | PBS102 | Thorlabs |
| Cylindrical lens | CL-1 | f = -75 mm | LK1432RM-A | Thorlabs |
| Cylindrical lens | CL-2 | f = 150 mm | LJ1629RM-A | Thorlabs |
| Cylindrical lens | CL-3 | f = 100 mm | LF1567RM-A | Thorlabs |
| Galvos system, 2-dimensional | GV | 6-mm aperture, silver mirrors | H8363 | Cambridge Tech. |
| Scan lens | SL | f = 110 mm | LSM05-BB | Thorlabs |
| Tube lens, illumination | TL-I | f = 200 mm | AC508-200-B | Thorlabs |
| Illumination objective, 10X | IO | NA = 0.3, f = 20 mm, water immer., WD = 3 mm | CFI Plan Fluor 10X W | Nikon |
| Sample chamber | SC | Machined from custom-design; black delrin | N/A | Proto Labs |
| Detection objective 16X | DO | NA = 0.8, f = 12.5 mm, water immer., WD = 3 mm | CFI75 LWD 16X W | Nikon |
| Detection objective 20X | DO | NA = 0.5, f = 9 mm, water immer., WD = 3.5 mm | UMPLFLN 20XW | Olympus |
| Filter, detection (green fluorescence) | FT | Pass band = 525/50 nm; 25-mm diameter | FF03-525/50-25 | Semrock |
| Filter, detection (red fluorescence) | FT | Pass band = 593/46 nm; 25-mm diameter | FF01-593/46-25 | Semrock |
| Tube lens, detection (used with 16X DO) | TL-D | f = 400 mm | AC508-400-A | Thorlabs |
| Tube lens, detection (used with 20X DO) | TL-D | f = 180 mm | AC508-180-A | Thorlabs |
| Micro-lens array | LA | Pitch = 150 μm, f = 3.0 mm, area = 16 x 16 mm<sup>2</sup> | APO-Q-P150-F3.0(633) | Flexible Optical B.V. |
| Relay lens | RL-1, RL-2 | f = 50 mm; f/1.4 (NIKKOR) |  | Nikon |
| Detection camera | DC | Scientific CMOS, pixel size = 6.5 μm, 2560 x 2160 pixels | Zylar 5.5 | Andor |

f: focal length
NA: numerical aperture

**Supplementary Table 2.** Imaging and reconstruction parameters for all presented results.

| Figures and Movies | Primary detection objective | Actual Mag. (X) | Mode (Excitation wavelength, nm) | Illumination z-extent ($\mu\text{m}$) | z-spacing ($\mu\text{m}$) | xy-pixel, acq. ($\mu\text{m}$) | xy-pixel, recon. ($\mu\text{m}$) | Detect. Band-pass (nm) |
| --- | --- | --- | --- | --- | --- | --- | --- | --- |
|  |  | (a) |  | (b) | (c) | (d) | (e) |  |
| Fig. 1c (SPIM) | 16X, NA = 0.8 | 32 | SPIM (488) | 3.5 | 1 | 0.203 | 0.311 | 525/50 |
| Fig. 1c (SVIM) | 16X, 0.8NA | 32 | SVIM (488) | 100 | 2 | 0.203 | 0.311 | 525/50 |
| Fig. 1d, Sup. Fig. 4a (SPIM) | 20X, NA = 0.5 | 20 | SPIM (488) | 3.5 | 1 | 0.325 | 0.498 | 525/50 |
| Fig. 1d (SVIM, 100 $\mu\text{m}$) | 20X, NA = 0.5 | 20 | SVIM (488) | 100 | 2 | 0.325 | 0.498 | 525/50 |
| Fig. 1d (SVIM, 300 $\mu\text{m}$) | 20X, NA = 0.5 | 20 | SVIM (488) | 300 | 2 | 0.325 | 0.498 | 525/50 |
| Fig. 1d, Sup. Fig. 4f (Wide-field) | 20X, NA = 0.5 | 20 | WF-LFM (488) | Entire sample | 2 | 0.325 | 0.498 | 525/50 |
| Fig. 2a | 20X, NA = 0.5 | 20 | WF-LFM (488) | Entire sample | n/a | 0.325 | 0.498 | 525/50 |
| Fig. 2b, c, d | 20X, NA = 0.5 | 20 | SVIM (488) | 100 | 2 | 0.325 | 0.498 | 525/50 |
| Fig. 2e-f (Blood) | 16X, NA = 0.8 | 32 | SVIM (561) | 150 | 2 | 0.203 | 0.311 | 593/46 |
| Fig. 2e-f (Endocardium) | 16X, NA = 0.8 | 32 | SVIM (488) | 150 | 2 | 0.203 | 0.311 | 525/50 |
| Fig. 3 (Wide-field ill.) | 20X, NA = 0.5 | 20 | WF-SVIM (488) | Entire sample | 2 | 0.325 | 0.498 | 525/50 |
| Fig. 3 (1-phonton ill.) | 20X, NA = 0.5 | 20 | 1p-SVIM (488) | 100 | 2 | 0.325 | 0.498 | 525/50 |
| Fig. 3 (2-photon ill.) | 20X, NA = 0.5 | 20 | 2p-SVIM (920) | 100 | 2 | 0.325 | 0.498 | 525/50 |
| Suppl. Fig. 2 | 16X, 0.8NA | 32 | SVIM (488) | 100 | 2 | 0.203 | 0.311 | 525/50 |
| Movie 1, Movie 2 | Rendered from the same results represented in Fig. 2c-d |  |  |  |  |  |  |  |
| Movie 3, Movie 4 | Rendered from the same 2-color data represented in Fig. 2e |  |  |  |  |  |  |  |

NA: Numerical aperture; WF: Wide-field; Mag.: Magnification.
(a) Actual magnification was determined by the ratio of the focal length of the primary detection objective over the focal length of the tube lens used.
(b) Illumination z-extent in SPIM mode was determined by the nominal thickness of the light sheet at the center of the field of view. For SVIM modes, the illumination z-extent was controlled by setting the scanning amplitude of the appropriate illumination galvo (see Methods).
(c) Spacing in z-direction for SPIM was determined by the z-step size of the acquired z-stack. Spacing in SVIM was computationally set at the onset of the light field reconstruction.
(d) Pixel size, acquired, was determined by the physical size of the camera pixel (6.5 $\mu\text{m}$ ) divided by the actual imaging magnification.
(e) Pixel size, reconstructed, was computationally set at the onset of the light field reconstruction.

**Supplementary Movie 1**

**Raw light-field-image movie of the bacterial flow around the squid light organ.**

Movie depicts the raw, unprocessed 2D-light field images of the flow of fluorescently-labeled *Vibrio fischeri* bacteria around the light organ of the Hawaiian bobtail squid *Euprymna scolopes*, during early stage of colonization. Light field images were acquired at 20 frames/s, and movie playback frame rate was set at the same rate. Note the high contrast and resolution achieved in these raw light field images, which enabled following the 3D motion of individual bacterium. A single frame of the movie was shown in Fig. 1b. Scale bar, 100 µm.

**Supplementary Movie 2**

**3D flow fields of bacteria around the squid light organ.**

Movie depicts the flow fields of fluorescently-labeled *Vibrio fischeri* bacteria around the light organ of the Hawaiian bobtail squid *Euprymna scolopes*, tracked from the 3D-reconstructed light field rendering. Light field images were acquired at 20 frames/s, yielding 3D volumetric rate of 20 volumes/s after reconstruction, of volume ∼ 600 x 600 x 100 (depth) µm^3^. Movie playback frame rate was set to 20 frames/s. Gray-scale image in the background is the average intensity projection of the 3D reconstruction. Individual fluorescent bacterium was computationally tracked, and tracks are shown color-coded by z position. Analysis of the tracks provided a quantitative description of the 3D bacterial flow fields, as shown in Fig. 2c,d. Scale bar, 100 µm.

**Supplementary Movie 3**

**3D blood flow and endocardium motion of the entire larval zebrafish beating heart.**

Movie of fluorescently-labeled endocardium (white) and blood cells (red) in a beating heart of a live 5-dpf zebrafish larva, 3D-rendered following the light field reconstruction. Acquisition was at 90 volumes/s, and movie playback frame rate was slowed by 5.5 times. Representative blood cells’ trajectories through the heart were manually tracked and quantified (color of the trajectories depict speed). During the movie, the endocardium channel was turned off at several timepoints to aid visualization of the blood cells. The high synchronous volumetric imaging rate and volume coverage enabled direct imaging and tracking of the quasi-chaotic blood flow, at single-blood-cell resolution throughout the cardiac beating cycle, which was about 450 ms. Still frames from the movie were depicted in Fig. 2e-h. Scale bar, 50 µm.

**Supplementary Movie 4**

**3D trajectories of individual blood cells flowing through the larval zebrafish beating heart.**

Movie depicts the representative trajectories of blood cells flowing through the larval zebrafish beating heart, as seen while rotating the imaged cardiac volume about the y-axis. Note the significant extent of the motion along the z direction, and the non-uniform 3D trajectories, of the blood cells. Acquisition was at 90 volumes/s, and movie playback frame rate was slowed by 5.5 times. Centroids of blood cells are shown as black-colored spheres. Each individual blood cell trajectory is represented in a single color to aid visualization. A 3D-cropped still frame of the movie was shown in Fig. 2h. Scale bar, 50 µm.

## References

1. Pawley, J. B. Handbook Of Biological Confocal Microscopy. (Springer-Verlag US, 2006).

2. Yang, W. & Yuste, R. *In vivo* imaging of neural activity. Nat. Methods 14, 349–359 (2017).

3. Levoy, M., Ng, R., Adams, A., Footer, M. & Horowitz, M. Light field microscopy. ACM Trans. Graph. TOG 25, 924–934 (2006).

4. Broxton, M. et al. Wave optics theory and 3-D deconvolution for the light field microscope. Opt. Express 21, 25418 (2013).

5. Prevedel, R. et al. Simultaneous whole-animal 3D imaging of neuronal activity using light-field microscopy. Nat. Methods 11, 727–730 (2014).

6. Huisken, J. Optical Sectioning Deep Inside Live Embryos by Selective Plane Illumination Microscopy. Science 305, 1007–1009 (2004).

7. Trivedi, V. et al. Dynamic structure and protein expression of the live embryonic heart captured by 2-photon light sheet microscopy and retrospective registration. Biomed. Opt. Express 6, 2056–2066 (2015).

8. Nyholm, S. V. & McFall-Ngai, M. The winnowing: establishing the squid–*vibrio* symbiosis. Nat. Rev. Microbiol. 2, 632–642 (2004).

9. McFall-Ngai, M. Divining the Essence of Symbiosis: Insights from the Squid-Vibrio Model. PLOS Biol. 12, e1001783 (2014).

10. Nyholm, S. V., Stabb, E. V., Ruby, E. G. & McFall-Ngai, M. J. Establishment of an animal–bacterial association: Recruiting symbiotic vibrios from the environment. Proc. Natl. Acad. Sci. U. S. A. 97, 10231–10235 (2000).

11. Nawroth, J. C. et al. Motile cilia create fluid-mechanical microhabitats for the active recruitment of the host microbiome. Proc. Natl. Acad. Sci. 114, 9510–9516 (2017).

12. Millikan, D. S. & Ruby, E. G. Alterations in Vibrio fischeri Motility Correlate with a Delay in Symbiosis Initiation and Are Associated with Additional Symbiotic Colonization Defects. Appl. Environ. Microbiol. 68, 2519–2528 (2002).

13. Millikan, D. S. & Ruby, E. G. Vibrio fischeri Flagellin A Is Essential for Normal Motility and for Symbiotic Competence during Initial Squid Light Organ Colonization. J. Bacteriol. 186, 4315–4325 (2004).

14. Stainier, D. Y. R. Zebrafish genetics and vertebrate heart formation. Nat. Rev. Genet. 2, 39–48 (2001).

15. Santhanakrishnan, A. & Miller, L. A. Fluid Dynamics of Heart Development. Cell Biochem. Biophys. 61, 1–22 (2011).

16. Liebling, M., Forouhar, A. S., Gharib, M., Fraser, S. E. & Dickinson, M. E. Four-dimensional cardiac imaging in living embryos via postacquisition synchronization of nongated slice sequences. J. Biomed. Opt. 10, 054001 (2005).

17. Mickoleit, M. et al. High-resolution reconstruction of the beating zebrafish heart. Nat. Methods 11, 919 (2014).

18. Taylor, J. M. et al. Real-time optical gating for three-dimensional beating heart imaging. J. Biomed. Opt. 16, 116021 (2011).

19. Fahrbach, F. O., Voigt, F. F., Schmid, B., Helmchen, F. & Huisken, J. Rapid 3D light-sheet microscopy with a tunable lens. Opt. Express 21, 21010–21026 (2013).

20. Weber, M. et al. Cell-accurate optical mapping across the entire developing heart. eLife 6, e28307 (2017).

21. Hove, J. R. et al. Intracardiac fluid forces are an essential epigenetic factor for embryonic cardiogenesis. Nature 421, 172–177 (2003).

22. Vermot, J. et al. Reversing blood flows act through klf2a to ensure normal valvulogenesis in the developing heart. PLoS Biol. 7, e1000246 (2009).

23. Steed, E. et al. klf2a couples mechanotransduction and zebrafish valve morphogenesis through fibronectin synthesis. Nat. Commun. 7, 11646 (2016).

24. Truong, T. V., Supatto, W., Koos, D. S., Choi, J. M. & Fraser, S. E. Deep and fast live imaging with two-photon scanned light-sheet microscopy. Nat. Methods 8, 757–760 (2011).

25. Wolf, S. et al. Whole-brain functional imaging with two-photon light-sheet microscopy. Nat. Methods 12, 379–380 (2015).

26. Mukamel, E. A., Nimmerjahn, A. & Schnitzer, M. J. Automated analysis of cellular signals from large-scale calcium imaging data. Neuron 63, 747–760 (2009).

27. Keller, P. J., Schmidt, A. D., Wittbrodt, J. & Stelzer, E. H. K. Reconstruction of zebrafish early embryonic development by scanned light sheet microscopy. Science 322, 1065–1069 (2008).

28. Wu, Y. et al. Inverted selective plane illumination microscopy (iSPIM) enables coupled cell identity lineaging and neurodevelopmental imaging in Caenorhabditis elegans. Proc. Natl. Acad. Sci. 108, 17708–17713 (2011).

29. Bouchard, M. B. et al. Swept confocally-aligned planar excitation (SCAPE) microscopy for high-speed volumetric imaging of behaving organisms. Nat. Photonics 9, 113–119 (2015).

30. Tomer, R. et al. SPED Light Sheet Microscopy: Fast Mapping of Biological System Structure and Function. Cell 163, 1796–1806 (2015).

31. Quirin, S. et al. Calcium imaging of neural circuits with extended depth-of-field light-sheet microscopy. Opt. Lett. 41, 855–858 (2016).

32. Liu, T.-L. et al. Observing the cell in its native state: Imaging subcellular dynamics in multicellular organisms. Science 360, eaaq1392 (2018).

33. Cong, L. et al. Rapid whole brain imaging of neural activity in freely behaving larval zebrafish (Danio rerio). eLife 6, e28158 (2017).

34. Pégard, N. C. et al. Compressive light-field microscopy for 3D neural activity recording. Optica 3, 517–524 (2016).

35. Aimon, S. et al. Fast whole brain imaging in adult Drosophila during response to stimuli and behavior. bioRxiv 033803 (2017). doi:10.1101/033803

36. Grosenick, L. M. et al. Identification Of Cellular-Activity Dynamics Across Large Tissue Volumes In The Mammalian Brain. bioRxiv 132688 (2017). doi:10.1101/132688

37. Nöbauer, T. et al. Video rate volumetric Ca2+ imaging across cortex using seeded iterative demixing (SID) microscopy. Nat. Methods 14, 811–818 (2017).

38. Skocek, O. et al. High-speed volumetric imaging of neuronal activity in freely moving rodents. Nat. Methods 1 (2018). doi:10.1038/s41592-018-0008-0

39. Piatkevich, K. D. et al. A robotic multidimensional directed evolution approach applied to fluorescent voltage reporters. Nat. Chem. Biol. 14, 352–360 (2018).

40. Platisa, J. & Pieribone, V. A. Genetically encoded fluorescent voltage indicators: are we there yet? Curr. Opin. Neurobiol. 50, 146–153 (2018).

41. Ghosh, K. K. et al. Miniaturized integration of a fluorescence microscope. Nat. Methods 8, 871–878 (2011).

42. Grover, D., Katsuki, T. & Greenspan, R. J. Flyception: imaging brain activity in freely walking fruit flies. Nat. Methods 13, 569–572 (2016).

43. Kim, D. H. et al. Pan-neuronal calcium imaging with cellular resolution in freely swimming zebrafish. Nat. Methods 14, 1107–1114 (2017).

44. Cohen, N. et al. Enhancing the performance of the light field microscope using wavefront coding. Opt. Express 22, 24817–24839 (2014).

45. Antipa, N. et al. DiffuserCam: lensless single-exposure 3D imaging. Optica 5, 1 (2018).

46. Sharpe, J. et al. Optical Projection Tomography as a Tool for 3D Microscopy and Gene Expression Studies. Science 296, 541–545 (2002).

47. Zheng, G., Horstmeyer, R. & Yang, C. Wide-field, high-resolution Fourier ptychographic microscopy. Nat. Photonics 7, 739–745 (2013).

48. Adams, J. K. et al. Single-frame 3D fluorescence microscopy with ultraminiature lensless FlatScope. Sci. Adv. 3, e1701548 (2017).

## Methods – References

49. Edelstein, A. D. et al. Advanced methods of microscope control using μManager software. J Biol Methods 1, (2014).

50. Troll, J. V. et al. Peptidoglycan induces loss of a nuclear peptidoglycan recognition protein during host tissue development in a beneficial animal-bacterial symbiosis. Cell. Microbiol. 11, 1114–1127 (2009).

51. Dunn, A. K., Millikan, D. S., Adin, D. M., Bose, J. L. & Stabb, E. V. New rfp- and pES213-Derived Tools for Analyzing Symbiotic Vibrio fischeri Reveal Patterns of Infection and lux Expression In Situ. Appl. Environ. Microbiol. 72, 802–810 (2006).

52. Schindelin, J. et al. Fiji: an open-source platform for biological-image analysis. Nat. Methods 9, 676–682 (2012).

53. Peli, E. Contrast in complex images. JOSA A 7, 2032–2040 (1990).

54. Bex, P. J. & Makous, W. Spatial frequency, phase, and the contrast of natural images. JOSA A 19, 1096–1106 (2002).

